# Quantitative prediction and measurement of Piezo’s membrane footprint

**DOI:** 10.1101/2022.06.28.497996

**Authors:** Christoph A. Haselwandter, Yusong R. Guo, Ziao Fu, Roderick MacKinnon

## Abstract

Piezo proteins are mechanosensitive ion channels that can locally curve the membrane into a dome shape (Y. R. Guo, R. MacKinnon, 2017). The curved shape of the Piezo dome is expected to deform the surrounding lipid bilayer membrane into a membrane footprint, which may serve to amplify Piezo’s sensitivity to applied forces (C. A. Haselwandter, R. MacKinnon, 2018). If Piezo proteins are embedded in lipid bilayer vesicles, the membrane shape deformations induced by the Piezo dome depend on the vesicle size. We employ here membrane elasticity theory to predict, with no free parameters, the shape of such Piezo vesicles outside the Piezo dome, and show that the predicted vesicle shapes agree quantitatively with the corresponding measured vesicle shapes obtained through cryo-electron tomography, for a range of vesicle sizes (Helfrich W. 1973). On this basis, we explore the coupling between Piezo and membrane shape, and demonstrate that the features of the Piezo dome affecting Piezo’s membrane footprint follow approximately a spherical cap geometry. Our work puts into place the foundation for deducing key elastic properties of the Piezo dome from membrane shape measurements and provides a general framework for quantifying how proteins deform bilayer membranes.

**Classification:** Biophysicss

## Main Text

### Introduction

In common with other organisms, vertebrates possess a variety of senses that respond to mechanical stimuli(1). Despite intense efforts, the molecules and physical mechanisms underlying vertebrate mechanosensation have long remained elusive. In 2010, Piezo proteins—Piezo 1 and Piezo 2 in mammals—were discovered, which has led to stunning progress in the elucidation of the molecular basis for mechanosensation(2). Piezo channels are mechanosensitive ion channels that open in response to mechanical force. They contain a central pore and three long arms that extend away from the protein center(3–5). The extended arms are made of transmembrane helices and do not lie in a plane in their closed conformation, which induces the lipid bilayer membrane to curve in-between the arms(3). We refer to the Piezo ion channel protein plus the curved lipid bilayer contained within the channel’s approximate perimeter as the ‘Piezo dome,’ with the lipid membrane connecting smoothly across the Piezo dome boundary. Piezo channels are thus expected to deform the lipid membrane outside the Piezo dome(6). We refer to the region of deformed lipid bilayer membrane outside the Piezo dome as Piezo’s ‘membrane footprint’(7). The mechanical properties of the membrane footprint may have interesting consequences for how Piezo responds to mechanical stimuli and may serve to amplify Piezo’s sensitivity.

The theoretical study of lipid bilayer-protein interactions has a long and rich history suggesting that protein-induced bilayer shape deformations can be captured by membrane elasticity theory(7–12). Membrane elasticity theory should therefore be able to account quantitatively for the membrane shape deformations induced by the Piezo dome. In particular, if Piezo proteins are embedded in lipid bilayer vesicles, membrane elasticity theory should be able to predict the shape of such Piezo vesicles outside the Piezo dome. The aim of this paper is three-fold: First, we provide quantitative comparisons between the predicted and observed shapes of Piezo vesicles. We thus test the continuum elasticity theory predicting the shape of Piezo’s membrane footprint, and develop a general framework for quantifying how proteins deform bilayer membranes. Second, we employ our experimental measurements and theoretical predictions of Piezo vesicle shape to learn about how Piezo curves the membrane. In particular, we investigate to what extent the geometry of the Piezo dome can be described by a spherical cap, as proposed previously based on cryo-electron microscopy (cryo-EM) and high-speed atomic force microscopy (HS-AFM) experiments(3, 13). Third, a quantitative understanding of Piezo vesicle shape can be used to provide insight into the elastic properties of the Piezo dome underlying its mechanosensory function, which we explore in a companion paper.

This study uses mathematical concepts and methods that might not be familiar to some readers. While the complete derivations for equations are presented in appendices, we still outline the mathematical concepts at each step in the main text. The paper is written, we hope, so that the logic can be followed and appreciated even if mastering some details might require further study(14–17).

## Results

### Measuring and parameterizing Piezo vesicles

Piezo 1 channels were reconstituted into POPC:DOPS:Cholesterol (8:1:1) vesicles, frozen in vitrified ice, and analyzed using cryo-EM tomography. Due to the intrinsic curvature of Piezo, the channels nearly exclusively reconstitute with their extracellular surface pointed towards the inside of the vesicle. For tomographic analysis, we identified Piezo vesicles containing Piezo near the outer edge of the vesicle, as viewed in projection in EM images (see Fig. 1*A*). Piezo vesicles are identifiable because the globular extracellular domain (see Fig. 1*A*) is visible adjacent to the membrane on the inside of the vesicle. Images were then collected while tilting the specimen stage between +42 and −42 degrees (see Fig. 1*B*). When three-dimensional (3D) tomographic reconstructions were generated, Piezo vesicles appeared to contain holes at opposite sides of the vesicle due to the limited range of tilt angles. Because we initially selected Piezo vesicles with Piezo near the outer edge of the vesicle as viewed in projection in EM images, we could define a cross-section of the vesicle that included Piezo and a band of membrane encircling the vesicle’s long axis. In other words, if we define Piezo’s location as the Piezo vesicle’s north pole, the tomogram defines a continuous strip of membrane around the Piezo vesicle that contains both the vesicle north and south poles, the south pole being that region on the vesicle surface furthest from Piezo (Fig. 1*B*). From this reconstruction, we cut out a thin plane that contains the long axis of the Piezo vesicle intersecting the north and south poles. This cut-out is called the oriented Piezo vesicle image. To be clear, the shape of the oriented Piezo vesicle image is generally not identical to the shape of the Piezo vesicle in projection in the sample image. The shapes would be the same if the Piezo channel were exactly on the edge of the Piezo vesicle in the projected EM image. The tomogram ensures that it is, within a small error, on the edge in the oriented image.

**Fig. 1.**
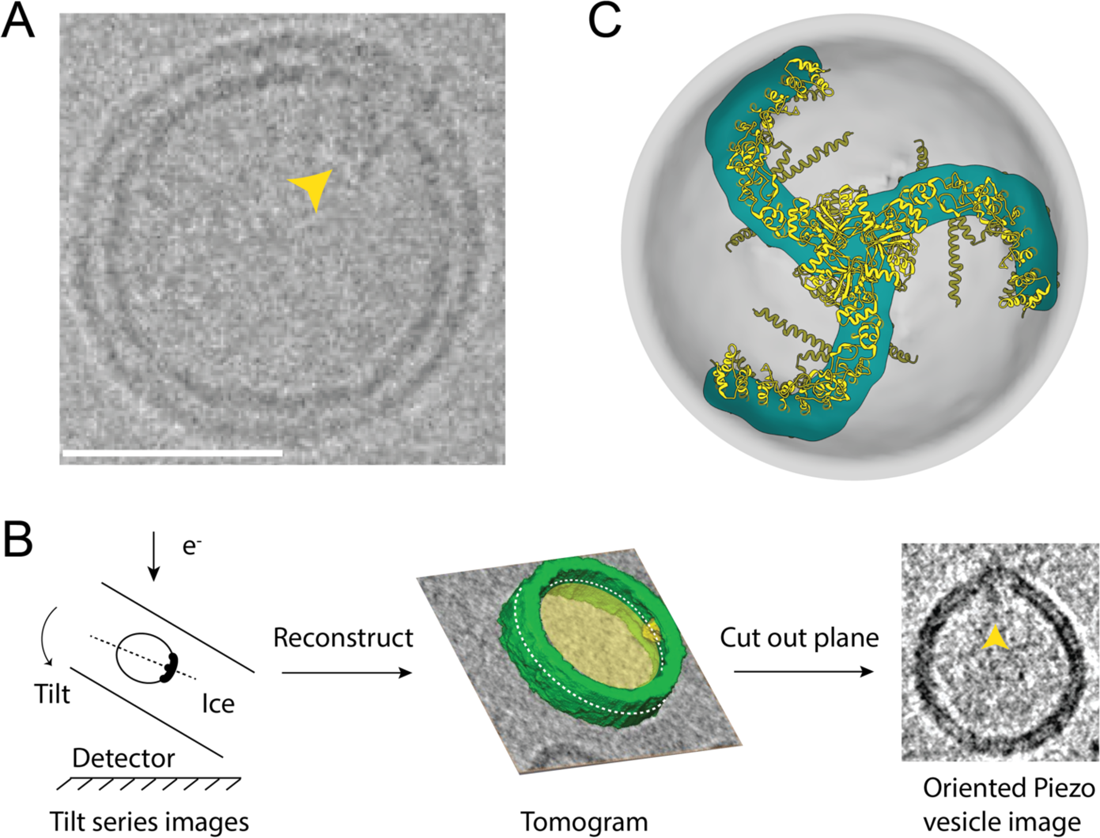
Measuring Piezo vesicles. (*A*) Cryo-EM image of a single Piezo vesicle with the Piezo cytoplasmic extracellular domain (CED) marked with a yellow pointer. The scale bar is 25 nm. (*B*) Schematic of Piezo vesicle data collection and tomographic image reconstruction. Left, tilt series images are collected on the detector while tilting the sample relative to the electron beam. Middle, a 3D tomogram is reconstructed from the tilt series images. The grey background and yellow interior correspond to one plane of the tomographic reconstruction. Part of the reconstructed vesicle is shown in green and the Piezo CED is colored yellow. The dashed white curve marks the intersection of the vesicle and a plane that includes the Piezo CED and the opposite “south” pole of the vesicle. Right, the oriented Piezo vesicle image shows the Piezo vesicle contour cut out from the intersecting plane. (*C*) Top-down view of the Piezo dome. The approximate position of the curved mid-bilayer surface of the Piezo dome is indicated in gray and its intersection with the Piezo protein in cyan. The ribbon model of Piezo1 (PDB: ID 6B3R) is colored yellow.

Piezo is a trimer of identical subunits with three-fold rotational symmetry (Fig. 1*C*). The lipids used in the present study form spherical vesicles in the absence of Piezo(3). Thus, we expect that within a short distance away from Piezo, excess free membrane folds will be minimized and the three-fold symmetry of the Piezo channel will give rise to ‘smooth-rotational symmetry’, i.e., order infinity, in the pure lipid membrane part of the vesicle. This assertion is corroborated by Piezo vesicles that appear circular in projection in EM images when viewed along Piezo’s three-fold axis. With this idea in mind, we generated the complete 3D shapes of Piezo vesicles from the oriented images as follows (see Fig. 2). The mid-membrane contour of oriented vesicle images was digitized manually and converted to a continuous, interpolated contour, as shown. The two sides of the interpolated contour, east and west, were reflected about the long vesicle axis connecting the vesicle north and south poles. The sides were averaged to produce a symmetrized Piezo vesicle profile, which we call the ‘Piezo vesicle profile’, and which defines the 3D surface of the Piezo vesicle through rotation about the central axis connecting the north and south poles. The border between the Piezo dome and the vesicle free membrane was defined by equating the surface area of the membrane region surrounding the vesicle north pole to the membrane surface area contained within Piezo’s perimeter, *A*_*P*_, estimated from the channel’s atomic coordinates (PDB ID 6B3R). This is the surface area of the Piezo dome, which includes the membrane-filled space in-between Piezo’s extended arms. To account for uncertainty—both due to inherent experimental limitations and the approximations used here—in defining the border separating the three-fold symmetric channel and the smooth-rotationally symmetric free membrane, in the accompanying paper, we consider variations in the Piezo dome area over the range 410 nm^2^ to 490 nm^2^ (<u>accompanying paper</u>). We call the surface area of the vesicle membrane outside the Piezo dome border *A*_*F*_, for ‘free’ membrane area. We specify the size of a vesicle using the variable *R*_*ν*_, the radius of a hypothetical sphere comprising the Piezo dome plus the free membrane: 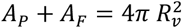.

**Fig. 2:**
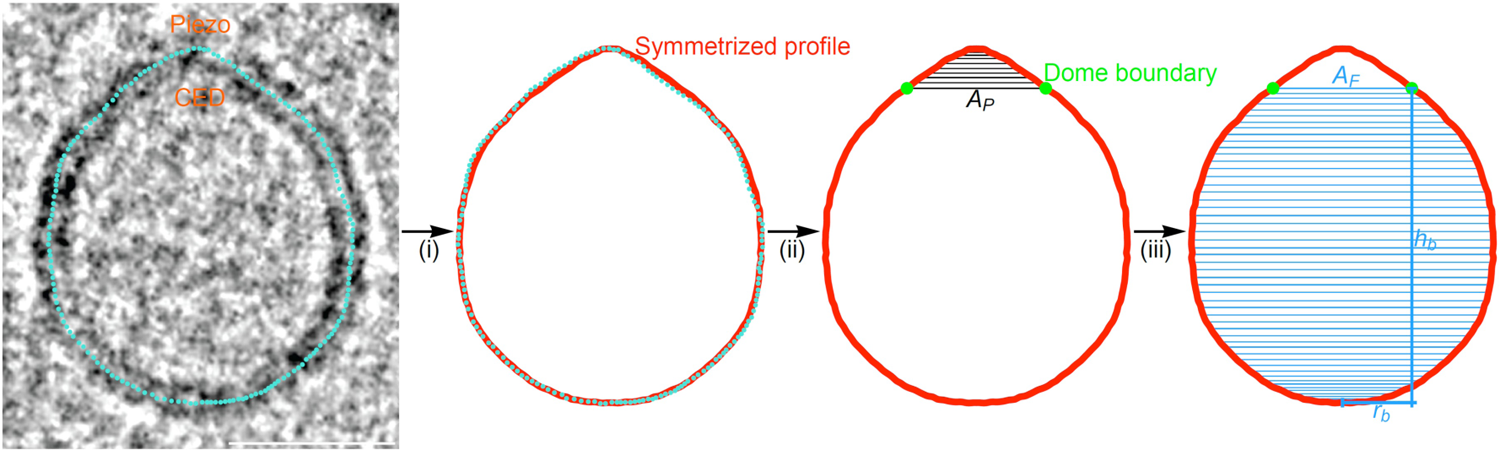
Quantifying Piezo vesicle profiles. Oriented Piezo vesicle image obtained by cryo-EM tomography (left panel). To quantify the Piezo vesicle shape, we manually trace the mid-membrane vesicle profile on the oriented Piezo vesicle image (cyan dots). Scale bar, 26 nm. (i) We take the line of maximal length from Piezo’s CED to the interpolated vesicle profile to correspond to the vesicle symmetry axis. Left-right averaging of the interpolated vesicle profile about the vesicle symmetry axis yields the symmetrized Piezo vesicle profile (red curve). (ii) We determine the Piezo dome boundary (green dots) by integrating out a vesicle surface area equal to the Piezo dome area, *A*_*P*_ ≈ 450 nm^2^, starting at the intersection of the vesicle symmetry axis and the Piezo dome (the vesicle north pole). (iii) As inputs for the membrane elasticity theory of Piezo vesicle shape, we extract from the measured (symmetrized) vesicle profiles the free vesicle area, *A*_*F*_, the in-plane radius of the Piezo dome, *r*_*b*_, and the height of the Piezo dome boundary above the vesicle south pole opposite the Piezo dome, *h*_*b*_. For a given Piezo vesicle radius *R*_*ν*_, *A*_*F*_ is determined by 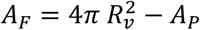, while the values of *r*_*b*_ and *h*_*b*_ follow directly from the observed positions of the Piezo dome boundary and of the vesicle south pole, respectively. (See also Figs. S1 and S2 in *Supplementary Information* section S1.)

Figures 3*A* and 3*B* show the 3D mid-bilayer surface of the free lipid membrane in an idealized Piezo vesicle and its profile on a graph with the vesicle south pole at the origin, respectively. We denote the arclength along the vesicle profile by *s*, with *s* = 0 at the south pole and *s* = *s*_*b*_ > 0 at the boundary where the free membrane meets the Piezo dome. In the analysis to follow, the free membrane region of a Piezo vesicle profile is represented parametrically with the horizontal *r*(*s*) and vertical *h*(*s*) coordinates given as functions of arclength, *s*, as shown in Fig. 3*B*. The coordinate system is thereby oriented so that *r*(*s*) corresponds to the radial distance from the long vesicle axis to a particular point on the vesicle surface, and *h*(*s*) corresponds to the height above the *r*-axis.

**Fig. 3:**
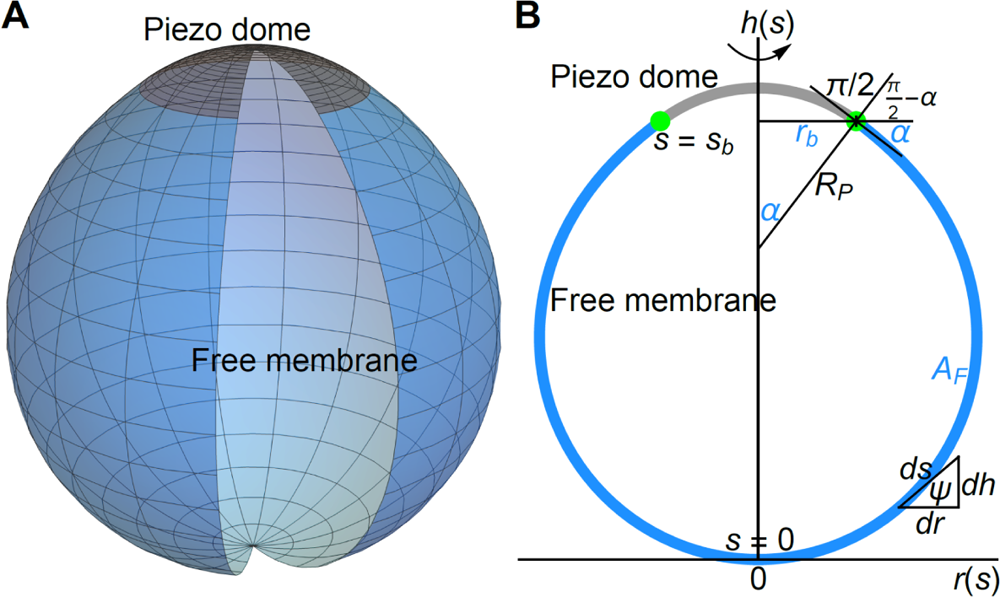
Membrane elasticity theory of Piezo vesicle shape. (*A*) 3D illustration of Piezo vesicles. The curved shape of the mid-membrane surface of the Piezo dome (indicated in grey) deforms the free lipid bilayer membrane in Piezo vesicles (indicated in blue). (*B*) Schematic of a Piezo vesicle profile in the vesicle symmetry plane containing Piezo’s central pore axis. We denote the arclength along the vesicle profile by *s*, with *s*_*b*_ = 0 at the vesicle south pole opposite the Piezo dome and *s* = *s*_*b*_ > 0 at the boundary (green dots) between the Piezo dome (grey curve) and the free lipid bilayer membrane outside the Piezo dome (blue curve). We represent the symmetry axis of the Piezo vesicle by the *h*-axis, the (in-plane) radial coordinate perpendicular to the *h*-axis by *r*, and the angle between the tangent to the vesicle profile and the *r*-axis (in the direction of increasing *r*) by *ψ* = *ψ*(*s*). For a given Piezo vesicle, the free membrane shape minimizing the Helfrich energy equation, Eq. 1, is determined by the values of three parameters, which are indicated in blue: The free membrane area *A*_*F*_, the in-plane Piezo dome radius *r*_*b*_, and the Piezo dome contact angle *α* = *π* − *ψ*(*s*_*b*_). We have the cord length (radius of curvature) 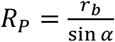 At the Piezo dome boundary. As illustrated by the triangle relating the differentials *ds, dr*, and *dh*, the shape variables satisfy the geometric relations 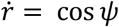 and 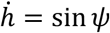.

### Strategy to predict free membrane shape

Next, we focus our attention on predicting the shape of the free membrane, that is, the part of the vesicle corresponding to 0 ≤ *s* ≤ *s*_*b*_ in Fig. 3*B*. You might wonder, if we can simply measure the shape experimentally, why would we want to predict it? The answer has two parts. First, our aim in this study is ultimately to deduce forces on Piezo from experimentally measured shapes of vesicles, which we pursue in a companion paper (<u>accompanying paper</u>). This is possible only if we understand why the Piezo vesicles are shaped the way they are, and prediction from theory is our working definition of understanding. Second, a membrane elasticity theory has been developed and tested over the past 50 years to successfully account for the experimentally observed shapes in pure lipid membrane systems such as giant unilamellar vesicles(18, 19). But in our case, we are studying frozen small unilamellar vesicles with a protein channel incorporated into them. Thus, we need to see how well the theory works in this circumstance.

Before presenting the mathematical approach to membrane shape prediction, we describe our basic strategy in words. We observe Piezo vesicles that contain one part protein and another part free lipid membrane. We have a theory for the ‘shape energy’ of free lipid membranes(20). The theory gives us an energy equation for the free membrane that lets us put in the free membrane shape and get out an energy. Note that we do not have such a theory for the protein. In fact, this is ultimately what we seek. So, for the free membrane part, how do we begin? We consider the protein as an object in the vesicle that exerts specific geometric constraints on the free membrane, i.e., boundary conditions, which we determine from experiment. Then, we focus our attention on the free membrane part of the vesicle and ask, given the boundary conditions and free membrane area, what free membrane shape is associated with the lowest energy? The core assumption here is that the lowest energy shape will correspond to the observed shape. In principle, one could try out all possible free membrane shapes that satisfy the boundary conditions by putting them into the energy equation and selecting the one that gives the lowest energy, but there are an infinite number of possible shapes. In practice, there is a mathematical approach, starting with the energy equation, the free membrane area, and the boundary conditions, to derive the lowest energy shape(14–16). This is what we mean by predicting the free membrane shape from the energy equation.

### The energy equation

The energy equation for the free membrane shape, due to Helfrich, is

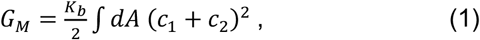

where the constant *K*_*b*_ is the lipid bilayer bending modulus and *c*_1_ and *c*_2_ are the principal curvatures of the mid-membrane surface, which are functions of position on the surface(20, 21). The integral is carried out over the entire free membrane vesicle surface. In words, every point on the free membrane surface is associated with a curvature number equal to the sum of the principal curvatures at that point. Equation 1 says that the elastic energy to bend the entire free membrane from a plane into its curved shape, *G*_*M*_, is computed by squaring the curvature number at each point, adding the values up, and multiplying the result by a constant, 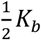. Because the lipids used in the present study do not produce membranes with discernable spontaneous curvature, Eq. 1 does not involve a term for ‘intrinsic’ curvature, i.e., its value is zero(20). Nor does it have a term for the Gaussian curvature, which is directly proportional to *c*_1_ *c*_2_, because, as mandated by the Gauss-Bonnet theorem, the Gaussian curvature contribution to Eq. 1 can be expressed as a fixed boundary term and a topological constant, and thus should not affect the free membrane vesicle shape(21). Also note, the Piezo vesicles were formed under conditions that preclude a pressure gradient across the membrane. Thus, vesicle volume is unconstrained, and the free membrane is under negligible tension.

To derive the lowest energy free membrane shape, that is, to minimize the energy equation, we follow the path developed for the study of axisymmetric vesicles(18, 22–24). We begin by expressing Eq. 1 using variables corresponding to the arclength parameterization of vesicle profiles (Fig. 3*B*). In addition to *r*(*s*) and *h*(*s*), for convenience we introduce a third variable, *ψ*(*s*), which is the angle between a line tangent to the vesicle profile and the *r*-axis (Fig. 3*B*). The variables *ψ*(*s*), *r*(*s*), and *h*(*s*), which we will call the shape variables, are related to each other through the geometric relations 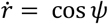 and 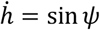; it is understood that *ψ, r*, and *h* are functions of the arclength parameter *s*, and the dot-notation means derivative with respect to *s* (Fig. 3*B*). Equation 1 then becomes

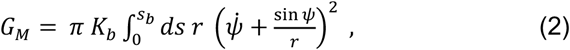

where 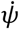 and 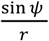 are the principal curvatures, *c*_1_ and *c*_2_, in the arclength parameterization. When minimizing Eq. 2, certain general conditions must be met. First, the geometric relations 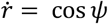 and 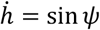 must be satisfied at every point on the free vesicle profile, and second, the total free membrane area must equal *A*_*F*_, the value of which will be determined from experiment. These conditions are enforced by introducing constraints into Eq. 2 through the mathematical method of Lagrange multipliers, to produce a constrained energy equation,

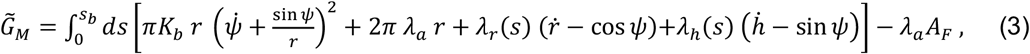

which is a function of the Lagrange multipliers *λ*_*r*_(*s*), *λ*_*h*_(*s*), and *λ*_*a*_ as well as the shape variables and (or) their derivatives with respect to *s*. The coefficients *λ*_*r*_(*s*) and *λ*_*h*_(*s*) in Eq. 3 associated with the geometric constraints depend on *s*, and *λ*_*a*_ is a constant coefficient for the term 2*π r*, which integrates to give the free membrane area and must satisfy 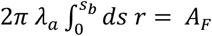. The utility of the Lagrange multipliers is this: the values, or functional forms, of the Lagrange multipliers are selected so that the constraints are met after Eq. 3 is minimized(15). But notably, when the constraints are met, i.e., when 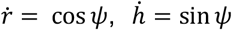, and 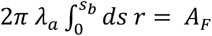, the terms multiplied by the coefficients vanish. This means that minimization of Eq. 3 will yield what we seek, the minimum of our original energy equation, Eq. 2 or Eq. 1, for a vesicle in which the constraints are satisfied.

### The Hamilton equations

For what comes next, we rewrite Eq. 3 as

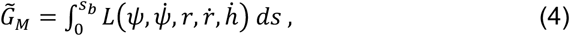

where *L*, called the Lagrangian function, stands for the integrand in Eq. 3 (16). If the right-hand side of Eq. 4 were an ordinary function, say *f*(*x,y*, …), at a minimum point its value would not change when *x* → *x* + *δx, y* → *y* + *δy*, … and we would find the values *x* = *x*_*min*_, *y* = *y*_*min*_, …associated with the minimum of *f*(*x,y*, …) by solving the derivative equations 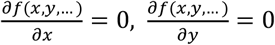 for *x, y*, … Similarly, the minimum of Eq. 4 occurs when small changes in the shape variables, i.e., *ψ* → *ψ* + *δψ, r* → *r* + *δr*, and *h* → *h* + *δh*, do not change the energy. By analogy to solving the derivative equations of an ordinary function, we could find the minimum energy form of the shape variables in Eq. 4 by solving ‘functional derivative’ equations called Euler-Lagrange equations 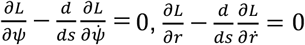, and 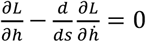, subject to specific boundary conditions at *s* = 0 and *s* = *s*_*b*_(16).

As a mathematically equivalent alternative to solving the Euler-Lagrange equations directly, we derive the minimum energy form of the shape variables by solving the Hamilton equations associated with Eq. 4(17). The Hamilton equations facilitate the numerical calculation of the free membrane shape of Piezo vesicles, and have the advantage that the boundary conditions encoding the key physical properties underlying Piezo vesicle shape can be imposed in a particularly transparent manner.

As outlined in the *Supplementary Information* section S2 appended to this article, the Hamilton equations associated with Eq. 4 are derived by switching from 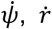 and 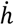 to 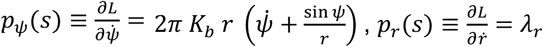, and 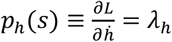 in the Euler-Lagrange equations through the Legendre transformation 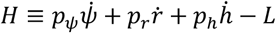, where *H* is known as the Hamiltonian(16, 17). *p*_*ψ*_ *p*_*r*_, and *p*_*h*_ are called generalized momenta, in reference to the physical meaning that these terms carry when describing dynamical systems in which the independent variable is time instead of arclength(16, 17). Following this Legendre transformation, rather than having one Euler-Lagrange equation for each shape variable *ψ, r*, and *h*, we have two first-order differential equations for each; one describing how the shape variable changes as a function of *s*, the other describing how its generalized momentum changes with *s*. The Hamilton equations associated with Eq. 4 are:

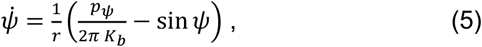

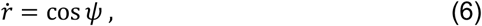

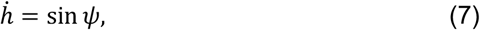

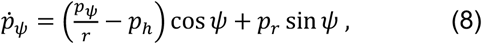

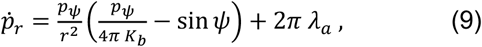

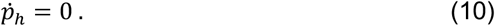

Note that the Lagrange multipliers *λ*_*r*_(*s*) and *λ*_*h*_(*s*) were eliminated in the derivation of the Hamilton equations, but that the associated geometric constraints on the free membrane are preserved in Eqs. 6 and 7. The Lagrange multiplier for the free membrane area, *λ*_*a*_, enters the Hamilton equations through Eq. 9; it is a parameter whose value must be determined from experiments. To apply the specific free membrane area constraint to each vesicle under study, we add to the system of Eqs. 5-10 a differential equation for the membrane area up to an arclength *s* from the south pole, *A*(*s*):

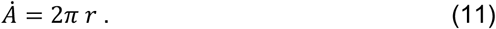

To predict the free membrane shapes of Piezo vesicles, we solve the 6 Hamilton equations, Eqs. 5-10, and the area equation, Eq. 11, subject to the boundary conditions at *s* = 0 and *s* = *s*_*b*_ associated with Piezo vesicles.

### Boundary conditions and input parameters

For a Piezo vesicle with the south pole at the origin of the *r-h* parametric plot we have, from geometry, *ψ*(*s*) = 0 and *r*(*s*) = 0 at *s* = 0. As shown in *Supplementary Information* section S3, *ψ*(0) = 0 and *r*(0) = 0 imply that *p* _*ν*_ (0) = 0. Since *h* does not appear explicitly in *L*, we are free to define *h*(0) = 0 (Fig. 3*B*). At *s* = *s*_*b*_, we assume that the free membrane and Piezo dome surfaces are tangent to each other, because the lipid membrane connects smoothly across the Piezo dome boundary. Note that, at *s* = *s*_*b*_, a cord of length 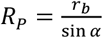, drawn perpendicular to the membrane so that it connects the long vesicle axis to the Piezo dome boundary, forms an angle *α* with the *h*-axis (Fig. 3*B*). We call *α* the contact angle. Therefore, from geometry, *ψ*(*s*_*b*_) = *π* − *α*. Furthermore, the projected, perimeter radius of the Piezo dome is given by *r*(*s*_*b*_) = *r*_*b*_ (Fig. 3*B*). Note that the values of *ψ*(*s*_*b*_) and *r*(*s*_*b*_) are fixed in a Piezo vesicle by the shape of the Piezo dome. By contrast, *h*(*s*_*b*_) is not fixed by Piezo because the length *h*(*s*_*b*_) − *h*(0) can freely vary when *G*_*M*_ is minimized, i.e., constrained physically by only the fixed values of *ψ*(*s*_*b*_), *r*(*s*_*b*_), and the free vesicle area, the vesicle could become shorter or longer, so its height is determined by the minimization of *G*_*M*_. Thus, we have what is called a zero-force or natural boundary condition on *h*(*s*) at *s* = *s*_*b*_ (14, 16). As shown in Appendix y, this means that *p*_*h*_(*s*_*b*_) = 0. But we already saw that 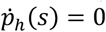, Eq. 10, which is to say *p*_*h*_(*s*) is unchanged over 0 ≤ *s* ≤ *s*_*b*_, and therefore, *p*_*h*_(0) = 0.

Another important aspect of the minimization of Eq. 4 is that the integration domain length, i.e., the arclength *s*_*b*_, is not fixed(22). Beginning at the south pole, the free membrane profile could take a circuitous path in getting to the edge of the Piezo dome as long as the boundary conditions already mentioned are satisfied and the free membrane area equals *A*_*F*_. As shown in *Supplementary Information* section S3, the variable integration domain length, together with the fact that *s* does not appear explicitly in *L*, leads to *p*_*r*_(0) = 0. For the membrane area we have *A*(0) = 0 and *A*(*s*_*b*_) = *A*_*F*_. Thus, in total, at *s* = 0 we have the seven ‘initial’ conditions

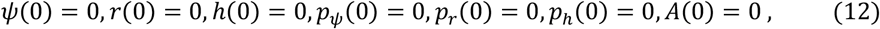

with which to solve the six Hamilton equations, Eqs. 5-10, and area equation, Eq. 11. However, these equations are associated with three parameters that are *a priori* unknown. We already saw that *λ*_*a*_, the Lagrange multiplier on the free membrane area, and *s*_*b*_, the upper limit of the integration domain, must be determined as part of the minimization of *G*_*M*_. In addition, the vesicle profile curvature at the south pole, 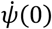, must be entered to solve Eqs. 5-11. To see this, note that the right-hand side of Eq. 5 evaluates to 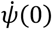 as *s* → 0, and thus 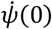 must be specified to make Eq. 5 meaningful at *s* = 0. While 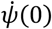 could in principle be estimated from the observed vesicle profiles, it is difficult to measure curvature accurately. For this reason, we set 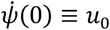 (23), with the value of *u*_0_, along with the values of *λ*_*a*_ and *s*_*b*_, to be determined from experimental constraints on the shape of each measured Piezo vesicle.

To solve the six Hamilton equations and area equation, Eqs. 5-11, associated with the three unknown parameters *λ*_*a*_, *s*_*b*_, and *u*_0_, we supplement the seven initial conditions in Eq. 12 with three endpoint boundary conditions measured from the Piezo vesicle profiles. As seen above, *A*(*s*_*b*_) must equal the measured free membrane area, *A*_*F*_, *r*(*s*_*b*_) must equal the measured projection radius of Piezo, *r*_*b*_, and *ψ*(*s*_*b*_) must equal *π* minus the Piezo dome contact angle, *α*. In practice, because obtaining the value of *α* involves taking the ratio of derivatives of the observed vesicle profile, we cannot measure *α* as accurately as a position on the vesicle profile. Thus, we use the measured position of *h*(*s*_*b*_) = *h*_*b*_ in place of *ψ*(*s*_*b*_) = *π* − *α* when predicting the free vesicle shapes. Using an experimental value for *h*(*s*_*b*_) in this way does not violate the zero-force boundary condition on *h*(*s*_*b*_) at *s* = *s*_*b*_, which led to the initial condition *p*_*h*_(0) = 0 in Eq. 12. Employing a so-called shooting method (see *Supplementary Information* section S4), we repeatedly solve Eqs. 5-11 subject to Eq. 12 to numerically adjust the values of *λ*_*a*_, *s*_*b*_, and *u*_0_ so as to satisfy the experimental constraints

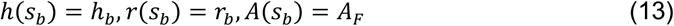

for each measured Piezo vesicle. Thus, for each Piezo vesicle we use the measured values of the three experimental input parameters *h*_*b*_ (in place of *α*), *r*_*b*_, and *A*_*F*_ to predict the free membrane shape through minimization of the Helfrich energy equation, without any free parameters.

### Free membrane shapes

Tomograms were collected on seven Piezo vesicles ranging in radius, *R*_*ν*_, from 12.1 nm to 36.2 nm. The vesicle profiles are shown in red (see Fig. 4). The dome boundary, in green, marks the point where the Piezo dome meets the free membrane, as defined by a Piezo dome area, *A*_*P*_ = 450 nm^2^. The coordinates of the dome boundary, *r*_*b*_ and *h*_*b*_, together with the free membrane area *A*_*F*_, were used to solve the Hamilton equations and area equation, Eqs. 5-11, subject to the initial conditions in Eq. 12 with the constraints in Eq. 13. The calculated free membrane profiles are shown in blue. The calculated profiles conform to the measured shapes of the Piezo vesicles and thus, the Helfrich energy equation appears to successfully predict the free membrane shape.

**Fig. 4:**
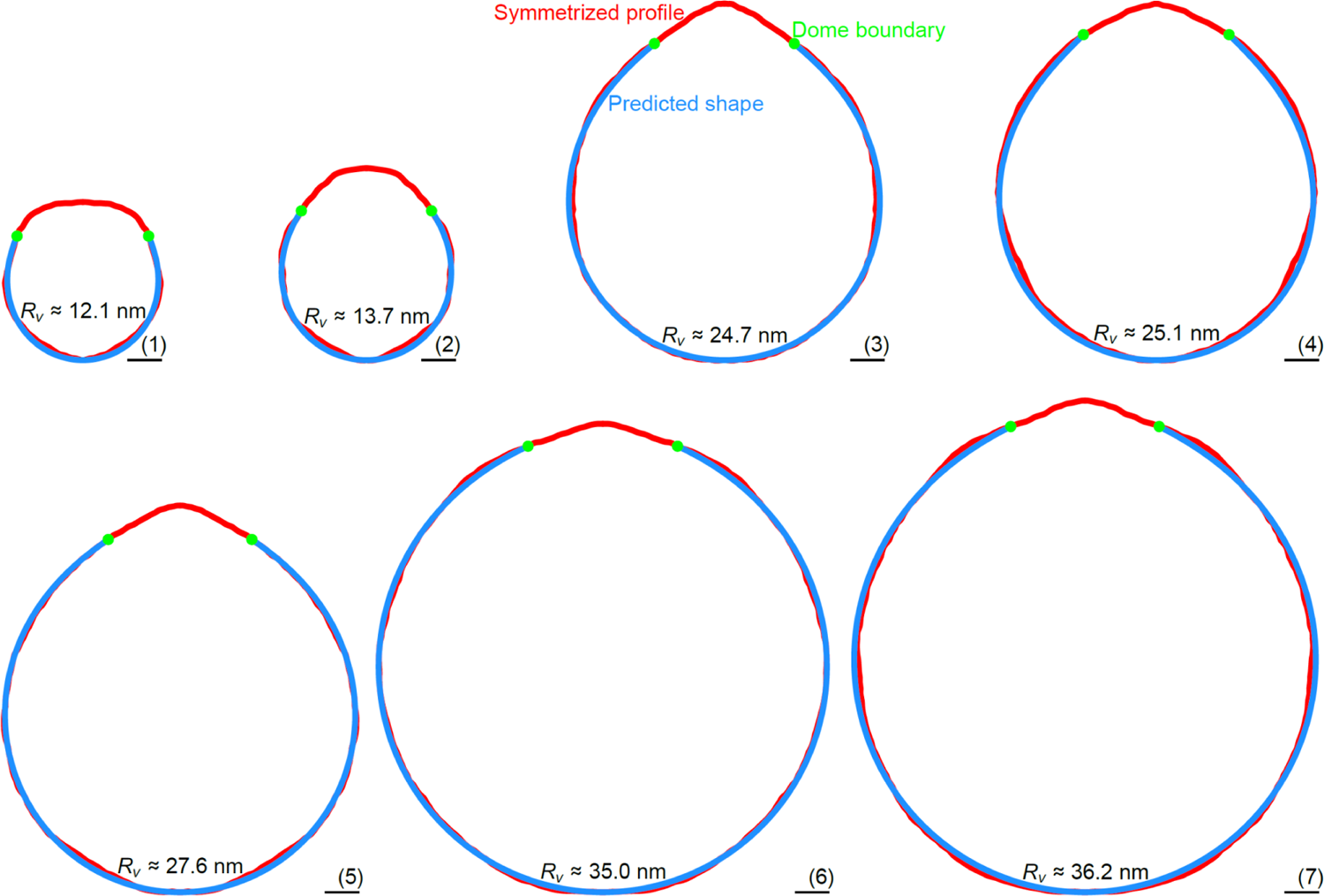
Quantitative prediction of Piezo vesicle shape. Symmetrized (measured) Piezo vesicle profiles (red curves) and predicted free membrane profiles (blue curves) for Piezo vesicles with radius (1) *R*_*ν*_ ≈ 12.1 nm, (2) *R*_*ν*_ ≈ 13.7 nm, (3) *R*_*ν*_ ≈ 24.7 nm, (4) *R*_*ν*_ ≈ 25.1 nm, (5) *R*_*ν*_ ≈ 27.6 nm, (6) *R*_*ν*_ ≈ 35.0 nm, and (7) *R*_*ν*_ ≈ 36.2 nm. The approximate root-mean-square deviations of the measured and predicted vesicle profiles are given by (1) 0.29 nm, (2) 0.35 nm, (3) 0.28 nm, (4) 0.49 nm, (5) 0.24 nm, (6) 0.25 nm, and (7) 0.44 nm. To calculate the blue curves we used, as inputs for the membrane elasticity theory of Piezo vesicle shape, the values of *A*_*P*_, *r*_*b*_, and *h*_*b*_ measured for each vesicle. The calculated free membrane profiles were obtained from the Helfrich energy equation, Eq. 1, without any free parameters. Scale bars, 5 nm. The oriented Piezo vesicle image in the left-most panel of Fig. 2 corresponds to vesicle 3.

There are several shortcomings to the approach used here that may limit the agreement between experimental and theoretical results. In particular, the predicted free vesicle shapes depend crucially on the measured values of *r*_*b*_, *h*_*b*_, and *A*_*F*_, which experiments can only yield approximately. Furthermore, the tomograms of Piezo vesicles do not produce a continuous vesicle surface around the long vesicle axis. As a result, we expect the measured, oriented Piezo vesicle images to be tilted slightly out of the planes defined by the vesicle cross sections, and hence to correspond only approximately to the predicted vesicle cross sections. Moreover, the location of the Piezo dome boundary can only be defined approximately in experiments, and the three-fold symmetry of the Piezo protein is expected to induce (slight) variations in the Piezo dome boundary conditions around the long vesicle axis. In contrast, our theoretical description of Piezo vesicles assumes perfect, smooth rotational symmetry of the free membrane shape. Considering these limitations, as well as the inevitable errors incurred when defining the mid-bilayer surface in the oriented Piezo vesicle images, the agreement between the predicted and measured free vesicle shapes in Fig. 4 appears to be remarkably good.

Up to this point, we have described the mathematical approach taken as one to find the minimum energy shape of the free membrane subject to geometric constraints arising from the Piezo dome shape and the vesicle size. In fact, the approach potentially identifies any shape that renders the free membrane energy unchanged to small perturbations of the shape variables(14, 16). This could include a local instead of a global minimum of the Helfrich energy equation, a maximum, or any other ‘stationary’ point. However, because our mathematical solutions match the experimentally observed shapes of Piezo vesicles that self-assembled under equilibrium conditions, it seems likely that the predicted free vesicle shapes indeed correspond to the global energy minimum of the free membrane shape.

Much consideration has been given to the concept that certain membrane proteins might distort the shape of their surrounding lipid bilayer membrane, creating a membrane footprint(6, 7, 9–12). Based on the membrane elasticity theory of bilayer-protein interactions, it has been proposed that the membrane footprint of proteins may affect membrane protein function, assist with membrane remodeling, yield bilayer-mediated protein interactions, and induce protein cooperativity. But it has been challenging to measure quantitatively and predict how proteins deform bilayer membranes, and hence to directly compare predicted and observed membrane footprints. The free membrane shapes of Piezo vesicles described here represent a direct measure of a protein’s membrane footprint, in the context of lipid bilayer vesicles, and show that membrane elasticity theory can be used to predict Piezo’s membrane footprint, thus linking membrane and protein shape.

### Piezo dome shapes

We next turn our attention to the parts of Piezo vesicles that contain Piezo. The Piezo dome in profile, shown in red between the green dome boundary marks (see Figs. 4 and 5*A*), has a somewhat irregular shape. This is not surprising since the Piezo dome contains the Piezo protein that is viewed perpendicular to its randomly oriented three-fold axis. Shape irregularities notwithstanding, systematic differences in the Piezo dome shape as a function of vesicle radius, *R*_*ν*_, are evident: in smaller vesicles the Piezo dome is more curved, and in larger vesicles it is flatter.

**Fig. 5:**
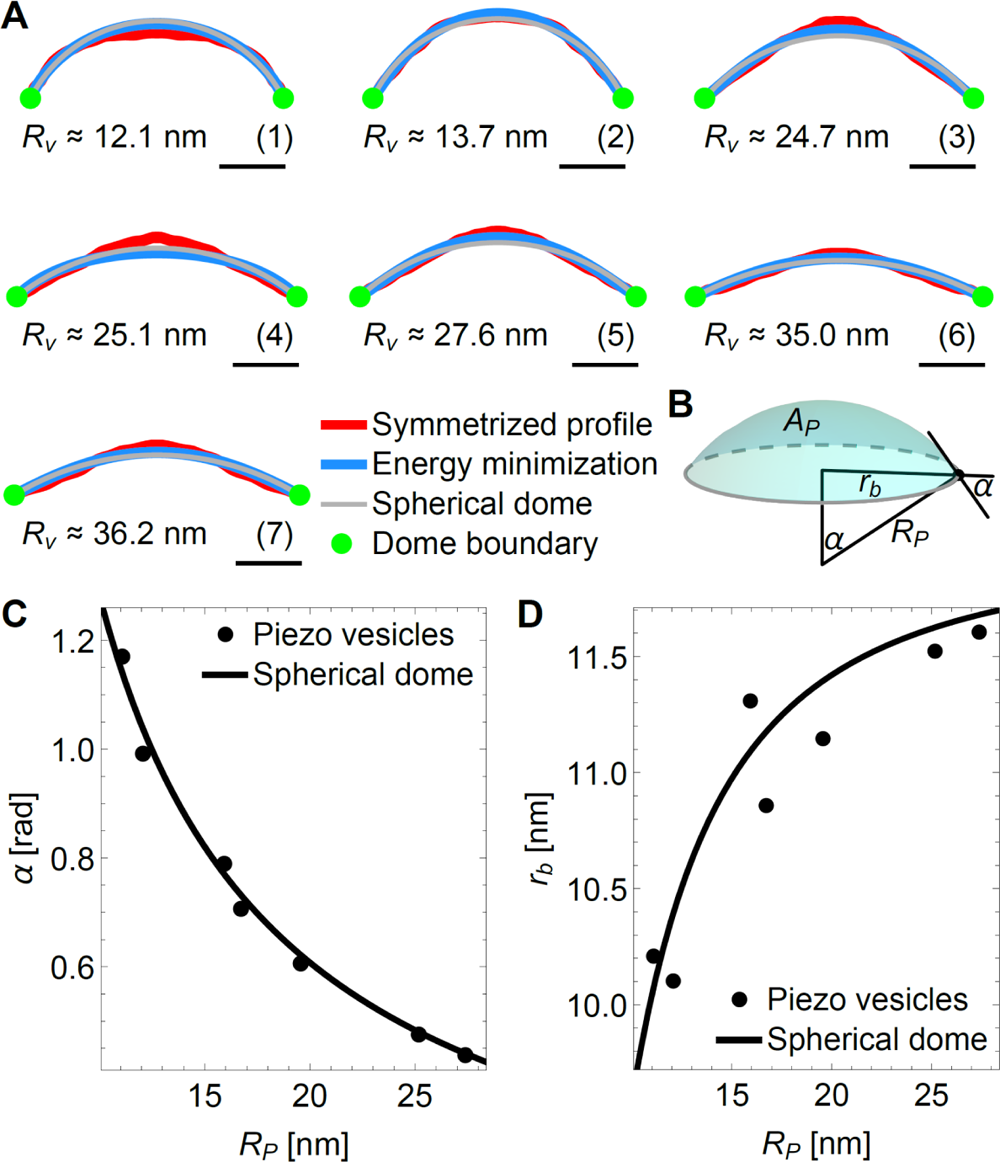
Coupling between Piezo and membrane shape. (*A*) Symmetrized (measured) Piezo dome profiles in Fig. 4 (red curves), Piezo dome shapes minimizing the Helfrich energy equation, Eq. 1, subject to *A*(*s*_*b*_) = *A*_*P*_ = 450 nm^2^ and the values of *α* and *r*_*b*_ associated with the predicted free membrane shapes in Fig. 4 (blue curves), and Piezo dome profiles corresponding to a spherical cap with area *A*_*CAP*_ = *A*_*P*_ and the in-plane radius *r*_*b*_ associated with the predicted free membrane shapes in Fig. 4 (grey curves). As in Fig. 4, the Piezo dome profiles correspond to Piezo vesicles with radius (1) *R*_*ν*_ ≈ 12.1 nm, (2) *R*_*ν*_ ≈ 13.7 nm, (3) *R*_*ν*_ ≈ 24.7 nm, (4) *R*_*ν*_ ≈ 25.1 nm, (5) *R*_*ν*_ ≈ 27.6 nm, (6) *R*_*P*_ ≈ 35.0 nm, and (7) *R*_*ν*_ ≈ 36.2 nm. Scale bars, 5 nm. (*B*) Schematic of the spherical cap model of the Piezo dome. The in-plane cap radius *r*_*b*_ and the cap angle *α* or, alternatively, the cap radius of curvature *R*_*P*_ and the cap area *A*_*CAP*_, completely define the geometric properties of the spherical cap. Piezo dome (*C*) contact angle *α* and (*D*) in-plane radius *r*_*b*_ versus radius of curvature *R*_*P*_ for the predicted free membrane shapes of the Piezo vesicles in Fig. 4 (points) and for a spherical dome (cap) with fixed area *A*_*CAP*_ = *A*_*P*_ (solid curves). For the Piezo vesicles, the values of *α* and *r*_*b*_ associated with the predicted free membrane shapes in Fig. 4 determine the radius of curvature at the Piezo dome boundary through 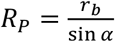 (see also Fig. 3*B*). For the spherical dome, the calculated curves follow from the geometric relations 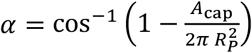 and 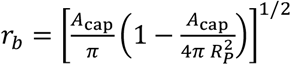 associated with spherical caps(25).

Is there a simple way to understand the observed Piezo dome shapes? We adapted the theory developed, above, for the free membrane outside the Piezo dome to the interior of the Piezo dome. The objective here is to determine whether the same physical principle governing the free membrane shape outside the Piezo dome—namely, minimization of the Helfrich energy equation— can also be used to describe the Piezo dome shape. Specifically, we minimized Eq. 1 through solution of Eqs. 5-11 with the initial conditions in Eq. 12 but now employing, as input parameter values, *A*(*s*_*b*_) = *A*_*P*_ and the values of *α* and *r*_*b*_ associated with the predicted free membrane shapes in Fig. 4. Note that *s* = 0 corresponds here to the vesicle north pole rather than the vesicle south pole. The resulting Piezo dome shapes, shown in blue, are seen to approximate the observed Piezo dome shapes (Fig. 5*A*). Curiously, these calculations suggest that the interior of the Piezo dome behaves somewhat like a flexible membrane, even though the Piezo dome contains the Piezo protein.

We also want to point out that the overall shape of the Piezo dome is not very far from a spherical cap (Fig. 5*A*). The shape of a spherical cap is conveniently specified through the cap radius of curvature *R*_*P*_ and the cap area *A*_CDE_(25). From the geometry of spherical caps we see that *R*_*P*_ and *A*_CDE_ can be used to define the Piezo dome properties affecting membrane shape deformations through 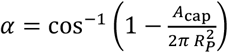 and 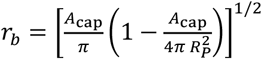 (see Fig. 5*B*). Previous cryo-EM and HS-AFM experiments suggest setting *A*_CDE_ = *A*_*P*_ in these equations, so that the shape of the Piezo dome is approximated by a spherical cap with fixed area *A*_*P*_ and variable radius of curvature *R*_*P*_ (3, 13). When we thus graph *α* and *r*_*b*_ as a function of *R*_*P*_ for the seven vesicles in Fig. 4, we find that *α* and *r*_*b*_ indeed vary as if the Piezo dome approximates a spherical cap of fixed area (Fig. 5*C*), suggesting that the features of the Piezo dome affecting Piezo’s membrane footprint can, approximately, be captured by a single parameter, the Piezo dome radius of curvature *R*_*P*_. We will use this result in our companion paper to study the mechanical response of the Piezo dome to the forces exerted by the surrounding lipid bilayer membrane (<u>accompanying paper</u>).

## Discussion

We have shown here that the Helfrich energy equation accurately predicts the observed shapes of the free membrane in Piezo vesicles over a range of vesicle sizes, without any free parameters. While it is not possible to directly measure Piezo’s membrane footprint in infinite, asymptotically flat membranes, our results suggest that membrane elasticity theory accurately captures protein-induced bilayer shape deformations from a length scale corresponding to molecular sizes to at least a length scale ∼00 nm.

Continuum elasticity theory implies that the free membrane shape of a particular Piezo vesicle follows from the values of three input parameters, which can all be estimated directly from experiments. In particular, we found here that the values of the free membrane area, *A*_*F*_, the projected radius of the Piezo dome, *r*_*b*_, and the height of the Piezo dome boundary above the opposite vesicle pole, *h*_*b*_, can be used to predict successfully the free vesicle shape. Other choices for the experimental input parameters could be used and give similar results.

Furthermore, we have shown here that Piezo changes its shape as a function of vesicle size, with the Piezo dome becoming less curved as the vesicle size is increased(13). We find that both the geometrical properties of the Piezo dome at its boundary and the shape of the Piezo dome conform approximately to a spherical cap with fixed area. Minimization of the membrane bending energy within the Piezo dome boundary also yields shapes close to the measured Piezo dome shapes. This seems surprising, given that the Piezo dome is not a uniform elastic membrane but contains the Piezo protein, and raises the fascinating prospect that the Piezo protein may be similarly flexible as the surrounding lipid bilayer membrane.

Instead of following the theoretical approach employed here, we could have substituted the observed shapes of the (smoothed) free vesicle membrane into the Helfrich energy equation, and hence deduced the membrane deformation energy associated with the observed free vesicle shapes. While this procedure would yield similar results for the membrane deformation energy, such an approach would be unable to predict the free membrane shape. Importantly, the agreement between the predicted and measured free membrane shapes suggests that the measured shapes correspond to minima of the Helfrich energy equation. With such a physical understanding of Piezo vesicle shape in hand, we can then ask: What would be the free membrane shape if the Piezo dome geometry were perturbed, and what would be the corresponding energy of the free membrane? As we show in a companion paper, the answer to this question allows calculation of the forces exerted by the Piezo dome on the surrounding lipid membrane and, conversely, on the Piezo dome by the surrounding lipid membrane (<u>accompanying paper</u>). From such a quantitative understanding of bilayer-protein interactions one can then deduce, based on the observed vesicle shapes, elastic properties of the Piezo dome underlying its mechanosensory function.

The simple version of the Helfrich energy equation employed here only involves a single term—the membrane bending energy—and only a single physical parameter—the bilayer bending rigidity, *K*_*b*_—that can be measured directly in experiments. The predicted free vesicle shapes are independent of the value of *K*_*b*_, and follow directly from minimization of the square of the sum of the two principal membrane curvatures across the free vesicle surface. The agreement between the predicted and measured free vesicle profiles suggests that, within experimental uncertainty, it is not necessary to account for effects other than those captured by minimization of the membrane bending energy, such as thermal fluctuations, bilayer coupling, or lipid tilt, to account for the observed Piezo vesicle shapes. Inclusion of such additional effects could further improve the agreement between observed and predicted vesicle shapes. Our results also suggest that the rapid freezing of Piezo vesicles for cryo-EM does not induce substantial perturbations of the free membrane shape. The agreement between predicted and measured free membrane shapes may thus be taken as an indication that cryo-EM can not only be used to accurately measure protein structures, but also to quantify the 3D shapes of lipid bilayers deformed by membrane proteins.

## Materials and Methods

### Proteoliposome sample preparation

Full-length mouse Piezo1 protein was expressed using the BacMam method as previously described(3). mPiezo1 was purified with a 6xHis tag at the C-terminus by cobalt affinity chromatography and size exclusion chromatography. Proteoliposomes (Piezo vesicles) with the lipid composition POPC:DOPS:cholesterol at a ratio of 8:1:1 (w:w:w) were prepared as previously described(13). To enrich the proteoliposomes for cryo-electron tomography, the liposome sample was subjected to cobalt affinity chromatography following reconstitution.

Freshly prepared proteoliposome samples were supplemented with 3 mM fluorinated fos-choline-8 (FFC-8) immediately before freezing. C-flat 200 mesh gold R1.2/1.3 holey carbon grids were glow-discharged prior to use. A first drop of 3.5 μl proteoliposome sample was applied to the grid, incubated for 15-60 s, and then manually removed using a filter paper. Then, a second drop of 3.5 μl proteoliposome sample was added, incubated for 15 s, and then blotted once for 1 s with −1 force and plunged into liquid ethane using a Vitrobot Mark IV (FEI company, Hillsboro, Oregon) operated at 22°C and 100% humidity. The grids were then stored in liquid nitrogen until imaging.

### Tilt-series data collection

Tilt-series were collected using a Titan Krios (Thermo Fisher) with a Gatan Bioquantum energy filter (Gatan) and a direct detector Gatan K3 (Gatan) at 300 keV. Data were collected symmetrically with a tilt range of −42° to 42° in 3° increments using SerialEM with 400-ms frames for each tilt image at a nominal defocus of 4 micrometers. The total dose per tilt-series collected was 103 e^−^ per Å^2^ with dose rates of approximately 20 e^−^ per pixel per s at a pixel size of 2.6 Å. Full-frame alignment was performed using MotionCor2(26).

### Tilt-series alignment

Tilt-series were aligned using Appion-Protomo(27–29). Tilt-series were coarsely aligned, manually aligned, and then refined using a set of alignment thicknesses. The best-aligned iteration was reconstructed for visual analysis using Tomo3D SIRT(30, 31) after dose compensation using a previously described method(32). CTF correction was not performed.

### Tomogram image processing

The reconstructed tomograms were visualized and analyzed in the 3DMOD. The coordinate of the Piezo1 CED in every liposome was determined by fitting the PDB model of mPiezo1 (PDB ID 6B3R). The distance between the CED and the outer membrane of the liposome was measured in the 3D reconstruction. The region of membrane associated with the longest distance away from the CED is defined as the south pole. The maximum projection plane (oriented Piezo vesicle image) was determined by drawing a plane intersecting the CED and the south pole.

The slice images of vesicles corresponding to the maximum projection plane were processed using an in-house script on the MATLAB R2021a platform (MathWorks) to annotate the mid-bilayer surface of the entire vesicle. Circles of diameter 5.7 nm, the average distance between phospholipid headgroup layers of our sample from previous measurements(13), were placed by visual inspection along the vesicle membrane defined by the headgroup layers. The (*x, y*) coordinates of the centers of these circles defined the mid-membrane contour of the vesicle within the maximum projection plane. A file of (*x, y*) coordinates was used to generate a vesicle profile using the *Interpolation*-command (with *InterpolationOrder* → 3) in *Mathematica(33)*.

### Prediction of Piezo vesicle shape

To predict the free membrane shapes of Piezo vesicles in Fig. 4 we numerically solved Eqs. 5-11 subject to Eqs. 12 and 13 using *Mathematica(33)*. We proceeded similarly in Fig. 5*A* to calculate the Piezo dome shapes through minimization of Eq. 1. A detailed description of mathematical derivations and methods can be found in the *Supplementary Information* sections S2-S4 appended to this article.

### Data availability

The traced Piezo vesicle profiles are provided in the file ‘Traced Piezo vesicle profiles.txt’ available online with this article. Figures S1 and S2 can be found in the *Supplementary Information* section S1 appended to this article. The tomograms of Piezo vesicles are deposited to EMDataBbank database with the accession codes D_1000265122, D_100025123, D_1000265124, D_1000265125 and D_1000265126.

## Acknowledgments

We thank Mark Ebrahim, Johanna Sotiris, and Honkit Ng at the Evelyn Gruss Lipper Cryo-EM Resource Center of Rockefeller University for assistance with cryo-EM data collection. C.A.H. thanks Jaime Agudo-Canalejo for useful correspondence regarding the shooting method in membrane mechanics. This work was supported at USC by NSF Grant No. DMR-2051681 and by NSF Grant No. DMR-1554716 (to CAH) and at Rockefeller University by NIH Grant GM43949 (to RM). R.M. is an investigator of the Howard Hughes Medical Institute.

